# Active and passive mechanics for rough terrain traversal in centipedes

**DOI:** 10.1101/2022.06.17.496557

**Authors:** Kelimar Diaz, Eva Erickson, Baxi Chong, Daniel Soto, Daniel I. Goldman

## Abstract

Centipedes coordinate body and limb flexion to generate propulsion. On flat solid surfaces, the limb-stepping patterns can be characterized according to the direction in which limb-aggregates propagate, opposite to (retrograde) or with the direction of motion (direct). It is unknown how limb and body dynamics are modified in terrain with terradynamic complexity more representative of their natural heterogeneous environments. Here, we investigated how centipedes that use retrograde and direct limp-stepping patterns, *S. polymorpha* and *S. sexspinosus*, respectively, coordinate their body and limbs to navigate laboratory environments which present footstep challenges and terrain rugosity. We recorded the kinematics and measured the locomotive performance of these animals traversing two rough terrains with randomly distributed step heights and compared the kinematics to those on a flat frictional surface. *S. polymorpha* exhibited similar body and limb dynamics across all terrains and a decrease in speed with increased terrain roughness. Unexpectedly, when placed in a rough terrain, *S. sexspinosus* changed the limb-stepping pattern from direct to retrograde. Further, for both species, traversal of rough terrains was facilitated by hypothesized passive mechanics: upon horizontal collision of a limb with a block, the limb passively bent and later continued the stepping pattern. While centipedes have many degrees of freedom. our results suggest these animals negotiate limb-substrate interactions and navigate complex terrains, by offloading complex control and leveraging the innate flexibility of their limbs.

## INTRODUCTION

How animals locomote and navigate their environments has been of interest to researchers and engineers (1), in inertia-dominated and non-inertial. In inertia-dominated systems, animals perform rapid locomotive behaviors and maneuvers (2; 3; 4; 5). Studies of these animals have led to the development of robot models capable of executing similar maneuvers (6; 7; 8). In contrast, in non-inertial systems, animals ranging from limbless to multi-legged must continuously self-deform to generate motion and overcome damping (9; 10; 11; 12; 13; 14).

Centipedes are animals with many limbs that are fast moving but are within the non-inertial regime (15). These animals locomote by generating and propagating a wave of limb flexion (termed here limb-stepping pattern) (15; 16). The limb-stepping pattern can be classified depending on the direction of propagation. When the limb-aggregates (i.e., grouped limbs) are propagated opposite to the direction of motion (of the animal) they are called *retrograde*, whereas when they are propagated with the direction of motion, they are called *direct* (17). Previously, Manton (16) characterized how distinct orders of centipedes use either direct or retrograde limb-stepping patterns. Centipedes of the order Scolopendramoprha, Geophilomorpha and Craterostigmorpha use retrograde limb-stepping patterns, while centipedes of the order Scutigeromorpha and Lithobiomorpha use direct limb-stepping patterns. Furthermore, centipedes that use retrograde or direct limb stepping patterns exhibit distinct body dynamics. Centipedes that use retrograde limb-stepping patterns exhibit body undulation, increasing body amplitude with increasing forward speed (16; 18). In contrast, centipedes that use direct limb-stepping pattern do not exhibit body undulation, even when stimulated to move at relatively high speeds (16; 18). However, what factors determine the selection of limb-stepping pattern remain unknown.

While Manton extensively researched centipedes and other arthropods (15; 16; 18), few studies in recent years have focused on centipedes’ locomotion (in part due to the difficulty to track many limbs). Previous studies have explored different aspects of centipede locomotion such as muscle activation patterns during body bending (19), gap traversal (20), effects of compromised appendages (i.e., missing limbs) (20), and effect of substrate friction (21). However, these studies have been limited to flat, solid, homogeneous terrains, unlike the animal’s natural environment. These animals can and must contend with heterogeneities (i.e., rocks, leaf litter, twigs) inherent to their ecology. In this regime, passive mechanics (e.g., limb flexibility) may be beneficial for locomotion on rugged terrains by reducing the complexity associated with precisely controlling the many degrees of freedom. Previous studies with other arthropods have shown that complex terrain traversal is achieved by passive mechanics such as passive mechanical response to specific events (i.e., adhesive pads in ants (22; 23)), and the distribution of passive mechanical elements (i.e., spines along limbs (14; 24)).

In contrast to few recent biological centipedes, synthetic (i.e., robots) multi-legged locomotors have become of interest over the last years. These have been developed to perform turning maneuvers (25; 26), navigate complex environments (27; 28; 29), overcome limb failures (30), among other capabilities. However, these were designed to serve as autonomous robots, few used as models (1; 31; 32) to explain centipede locomotive behaviors.

Here, we present the first study of biological centipedes locomoting on laboratory rough terrain for two species, *S. polymorpha* and *S. sexspinosus* (1A,B). These two centipede species have distinct kinematics on flat solid substrates; *S. polymorpha* uses retrograde limb-stepping pattern, whereas *S. sexspinosus* uses direct (1C,D). We study how these animals locomote complex terrain and what navigation strategies they use. We report the performance of these animals and found that *S. polymorpha* does not change locomotive strategy on complex terrain. In contrast, *S. sexspinosus* exhibits a change from direct to retrograde limb-stepping patterns. Further, we discovered an emergent passive behavior during limb-terrain interactions for both centipede species; a limb passively bent in the direction the force from the block was applied on it. Finally, we discuss the implications of gait switching in *S. sexspinosus* and possible advantages to the observed passive mechanics in both centipede species.

## MATERIALS AND METHODS

### Animals

All centipedes were wild caught. *S. polymorpha* were caught in Del Rio Val Verde County Texas. *S. sexspinosus* were caught in Valley National Park (CVNP), Summit County, Ohio. Four centipedes of each species were used in experiments with a mean body length of 7.7 ±1.5 cm and 6.2± 1.1 cm, for *S. polymorpha* and *S. sexspinosus*, respectively. *S. polymorpha* had 19 body segments with 19 joints and leg pairs. *S. sexspinosus* had 21 body segments with 21 joints and leg pairs. Centipedes were housed separately in plastic containers on a 12 hr:12 hr L:D photoperiod at room temperature (20-22°C). Centipedes were provided a source of water and fed mealworms weekly.

### Flat and rough terrains

Experiments were conducted on three different terrains (flat, lower rough, higher rough) placed in a glass tank (length = 51 cm, width = 27 cm, height = 32 cm) (Figure 2A). The flat terrain was a homogeneous level foamcore sheet. Rough terrains consisted of Gaussian (11; 27) and inverted Gaussian distributed (34) blocks of varying heights, for the lower and higher rough terrain, respectively (Figure 2B). Dimensions of each rough terrain was scaled to the size of each species body (length = 24 cm, width = 12 cm, for *S. polymorpha*; length = 16. width = 8 cm, for *S. sexspinosus*). Each rough terrain consisted of a 3D printed (Stratasys uPrint SE plus, material: ABSplus P430) height field, with 8 rows by 16 columns of square blocks (length and width = 1.5 cm for *S. polymorpha*, length and width = 1 cm for *S. sexspinosus*). For *S. polymorpha*, block heights varied from 0 to 1 cm and 0 to 1.5 cm, for the lower and higher rough terrain, respectively. For *S. sexspinosus*, block heights varied from 0 to 0.75 cm and 0 to 1 cm, for the lower and higher rough terrain, respectively. Terrains were placed level in the glass tank. All experiments were conducted at room temperature (20-22°C).

**Fig. 1.**
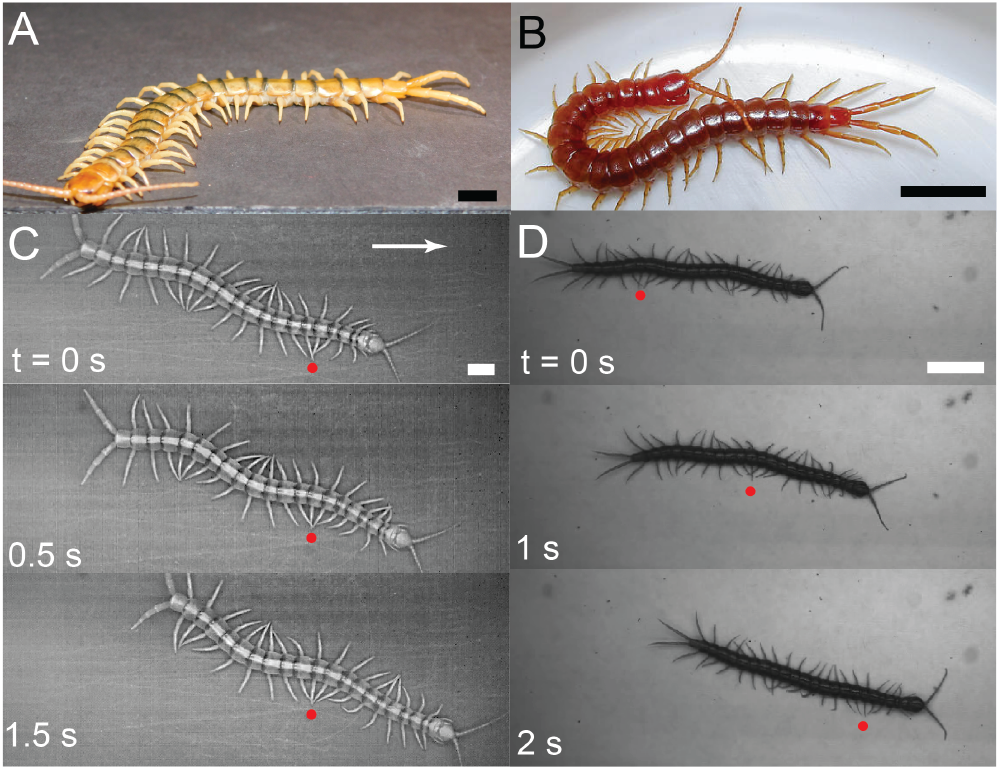
Centipedes with distinct limb-stepping patterns. Photo of (A) *Scolopendra polymorpha* and (B) *Scolopocryptops sexspinosus* (Image credit: Derek Hennen). Image sequence showing (C) *S. polymorpha* and (D) *S. sexspinosus* running on foam core. Red dots highlight a single location where adjacent limbs are aggregated. All scalebars correspond to 1 cm.

**Fig. 2.**
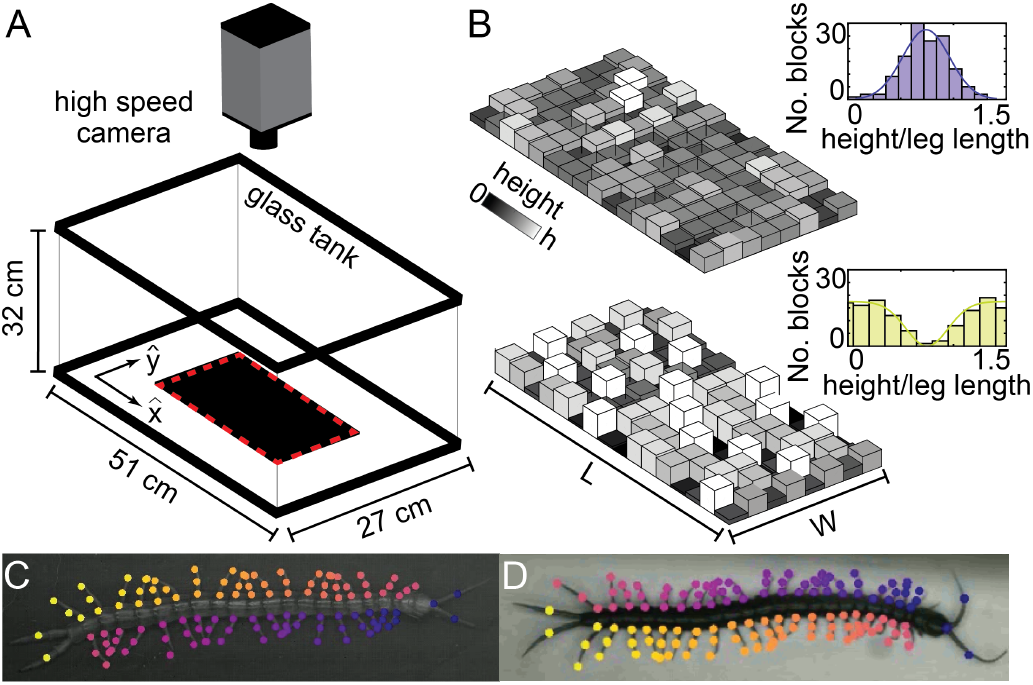
Experimental design and complex terrains. (A) Experimental set-up. Experiments were conducted in a 27 cm x 51 cm x 32 cm glass tank with a high speed camera placed vertically over the selected terrain. (B) Lower (top) rough terrain with Gaussian distributed blocks of varying heights. Inset shows the Gaussian distribution. Higher (bottom) rough terrain with inverted Gaussian distributed blocks of varying heights. Insets shows the Gaussian and inverted Gaussian distribution for the lower and higher rough terrain, respectively. Lower rough terrain block heights vary from 0 to 1 cm and from 0 to 0.75 cm for *S. polymorpha* and *S. sexspinosus*, respectively. Higher rough terrain block heights vary from from 0 to 1.5 cm and from 0 to 1 cm for *S. polymorpha* and *S. sexspinosus*, respectively. Blocks are colored by relative height. Labeled frame to track the kinematics of (C) *S. polymorpha* and (D) *S. sexspinosus* using DeepLabCut (33).

### Kinematic recordings

All experiments were recorded by a high speed camera (AOS, S-motion) positioned directly over the terrains to capture kinematics from a top-down view (Figure 2A). Videos were recorded at a resolution of 1280×700 pixels and a frame rate of 738 frames per second (fps). For both species, five videos per centipede (N = 4) per terrain were collected, with the exception of trials on the higher rough terrain for *S. polymorpha* for which a centipede lost a limb and died shortly after.

### Motion tracking

Positional data was extracted from videos with animal pose estimation software DeepLabCut (DLC) (33). Twenty frames from each video were manually labeled and then DLC provided positions for labeled points on all other frames. One hundred and thirty points per frame and 118 points per frame were labeled on the *S. polymorpha* and *S. sexspinosus*, respectively, including three points perlimb: body-limb contact, joint and tip, as well as points on the posterior and anterior antennae (Figure 2C,D). Points were placed within 0.5 cm of each limb position. Positional data obtained for *S. polymorpha* had a likelihood of 0.99 ±0.03, 0.98±.05, and 0.96±0.08 for flat, lower rough, and higher rough terrain, respectively. Positional data obtained for *S. sexspinosus* had a likelihood of 0.96±0.08, 0.95±0.15, and 0.94±0.12 for flat, lower rough, and higher rough terrain, respectively.

### Body and limb parameters

Digitized kinematics were used to calculate the body and limb parameters using custom MATLAB code. First, a Gaussian filter was used to smooth the x and y-coordinates of each tracked point. Filtered points of body-limb contact of opposite sides (left and right) were averaged over time to obtain a body midline. Body angles (*γ*) were obtained by finding the angle between the local tangent unit vector and the average direction of motion. Leg angles (*θ*) were obtained by calculating the angle between a limb and the local body normal (Figure 3A(i)).

**Fig. 3.**
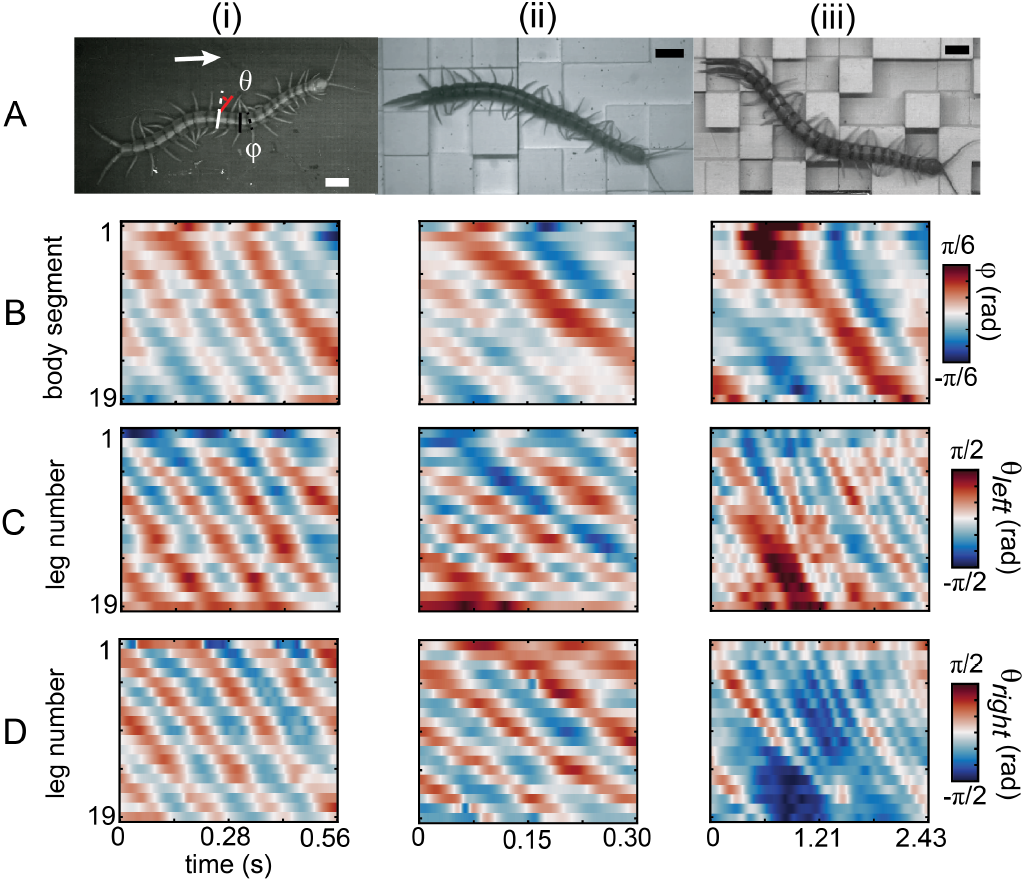
*S. polymorpha* locomoting on flat, lower rough, and higher rough terrain. (A) Images of *S. polymorpha* on (i) a flat surface, (ii) the lower rough terrain, and (iii) the higher rough terrain. Arrow shows the direction of motion of the animal (left to right) for all terrains. Body angles (*γ*) were obtained by finding the angle between the local tangent unit vector (black solid line) and the average direction of motion (black dashed line). Local tangent unit vector and average direction of motion are shifted from body midpoint to facilitate visualization. Leg angles (*θ*) were obtained by calculating the angle between a limb (dashed white line) and the local body normal (solid white line). Red circle corresponds to reconstructed body midpoint. All scalebars correspond to 1 cm. Heat maps show the generated by each (B) body segment and limb on both the (C) left and (D) right side of the animal over time.

Leg angles were used to calculate phase over time for each limb on the centipede’s body, similar to methods in (21). Each leg phase (*φ*_*i*_) was obtained from the difference between changes in the leg angle (Δ*θ*_*i*_) from the time average and its derivative 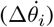 (Figure 6) (21). The retraction (*T*_*ret*_) period was obtained by finding the timing between *φ*_*i*_ = 0 and *φ*_*i*_ = *π*. The stride (*T*_*stride*_) period was obtained by calculating the timing between successive points when *φ*_*i*_ = 0. Duty factor (*DF*) was calculated as the ratio of the retraction and stride period (*DF* = *T*_*ret*_*/T*_*stride*_). Stride frequency was calculated as the inverse of the stride period (*ω*_*stride*_ = 1*/T*_*stride*_). Step length (*L*_*step*_) was obtained by calculating the total distance traveled for each associated *T*_*ret*_. Stride length (*L*_*stride*_) was calculated as the ratio of the step length and the study factor (*L*_*stride*_ = *L*_*step*_*/DF*). Because steps and/or stride could be interrupted due to limb-substrate collisions, *DF, ω*_*stride*_, *L*_*step*_, and *L*_*stride*_ were averaged for all limbs and the entirety of each trial. Statistical tests performed (t-test) for experimentally obtained parameters were performed using a custom MATLAB code.

## RESULTS

### Centipede kinematics

For both centipede species, we performed 20 trials on flat terrain, 20 trials on lower rough terrain, and 20 trials on higher rough terrain (16 in the case of *S. polymorpha*) (Figure 3, 4) (SI Movie 1). On the flat terrain, *S. polymorpha* exhibited body and limb-stepping waves propagated from head to tail (opposite to the direction of motion) along the body axis (Figure 3A-D(i)). In contrast, on the lower and higher rough terrain we observed the same limb-stepping pattern but no regular body undulation. With increasing terrain complexity, we observed interruptions in the limb-stepping patterns (Figure 3C-D(iii)). These correspond to limb-substrate collisions; as the centipede moves across the terrain, limb-substrate contact on the horizontal plane (i.e., limb contacting the side of a block) can occur due to the height disparities.

**Fig. 4.**
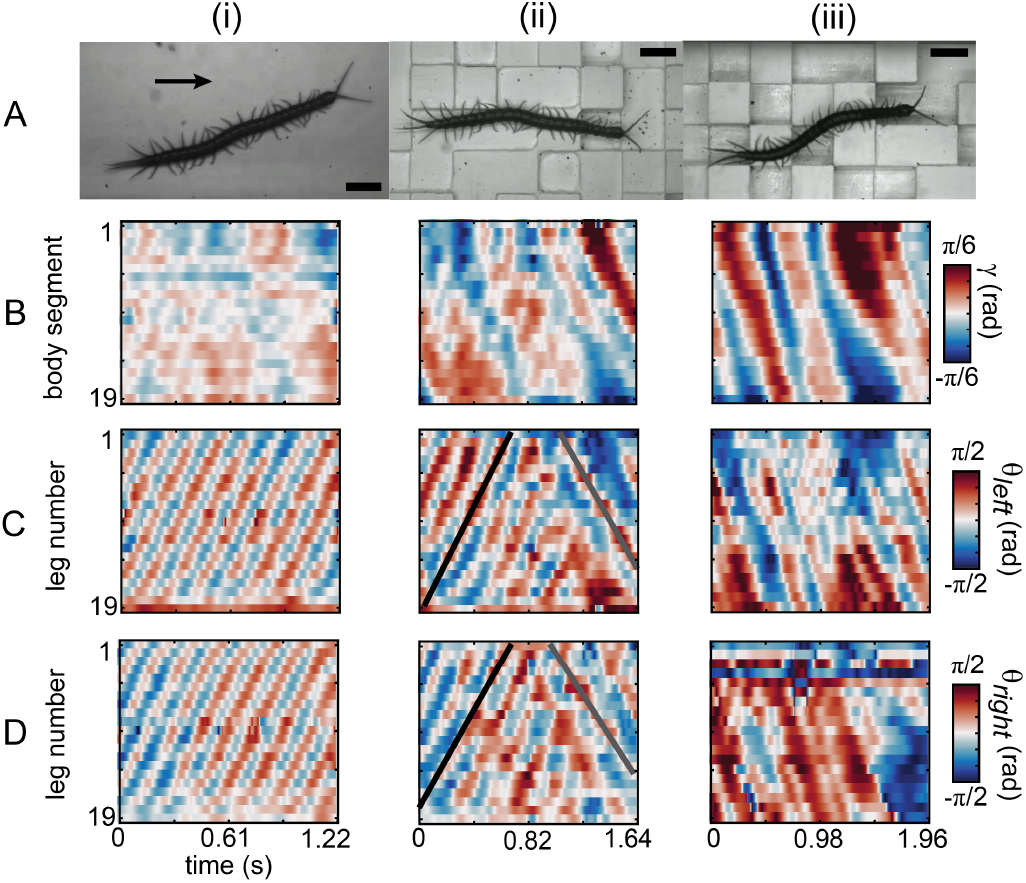
*S. sexspinosus* locomoting on flat, lower rough, and higher rough terrain. (A) Images of *S. sexspinosus* on (i) a flat surface, (ii) the lower rough terrain, and (iii) the higher rough terrain. Arrow shows the direction of motion of the animal (left to right). All scalebars correspond to 1 cm. Heat maps show the generated by each (B) body segment and limb on both the (C) left and (D) right side of the animal over time. Black line highlights when the animal used a direct limb-stepping pattern. Gray line highlights when the animal used a retrograde limb-stepping pattern.

On flat terrain, *S. sexspinosus* did not exhibit body undulation and propagated limb-stepping waves from tail to head (with the direction of motion) (Figure 4B-D(i)). This was surprising, as centipedes in the order Scolopendromorpha are thought to only use retrograde limb-stepping waves (16). On the lower rough terrain, *S. sexspinosus* demonstrated changes in their behaviors. On initial trials, the animals used solely direct limb-stepping waves. As more trials were collected, *S. sexspinosus* started the course using direct limb waves and changed the direction the limb wave was propagated from direct to retrograde (Figure 4C(ii),D(ii)). We note that, over time (i.e., minutes, from trial to trial), these centipedes would switch faster (i.e., earlier in the trial) from direct to retrograde or would only use retrograde limb-stepping waves. We hypothesize that the centipedes learned what terrain they locomoted on and actively changed the locomotive strategy. On the higher rough terrain, *S. sexspinosus* only used retrograde limb-stepping waves (Figure 4C(ii),D(ii)). Interestingly, we noticed a significant body undulation in *S. sexspinosus* on higher rough terrain; however, we posited that these body undulation is the passive response to the structures of rough terrain. Bands of body curvature observed in experiments (Figure 4B(ii),B(iii)) correspond to bends generated by the centipedes that are related to the path of travel through the terrain. Further, we observed similar interruptions in the limb-stepping pattern, relating to limb-substrate collisions in the horizontal plane due to height disparities between adjacent blocks (Figure 4C(iii),D(iii)).

### Performance across terrains

The direction in which the limbs propagate can be characterized by the leg phase shift (LPS). LPS is defined as the fraction of time during a gait cycle in which the forelimb leads the hindlimb in a set of two adjacent limbs (Figure 5A). A LPS < 0.5 corresponds to direct limb-stepping waves (propagated in the same direction of motion). In contrast, LPS > 0.5 corresponds to retrograde limbstepping waves (opposite to the direction of motion). A LPS = 0.5 corresponds to an alternating tripod gait. In hexapods, three leg pairs alternate ground contact forming a tripod. In myriapods, every other leg (e.g., all even numbered legs) in the same side has the same phase.

**Fig. 5.**
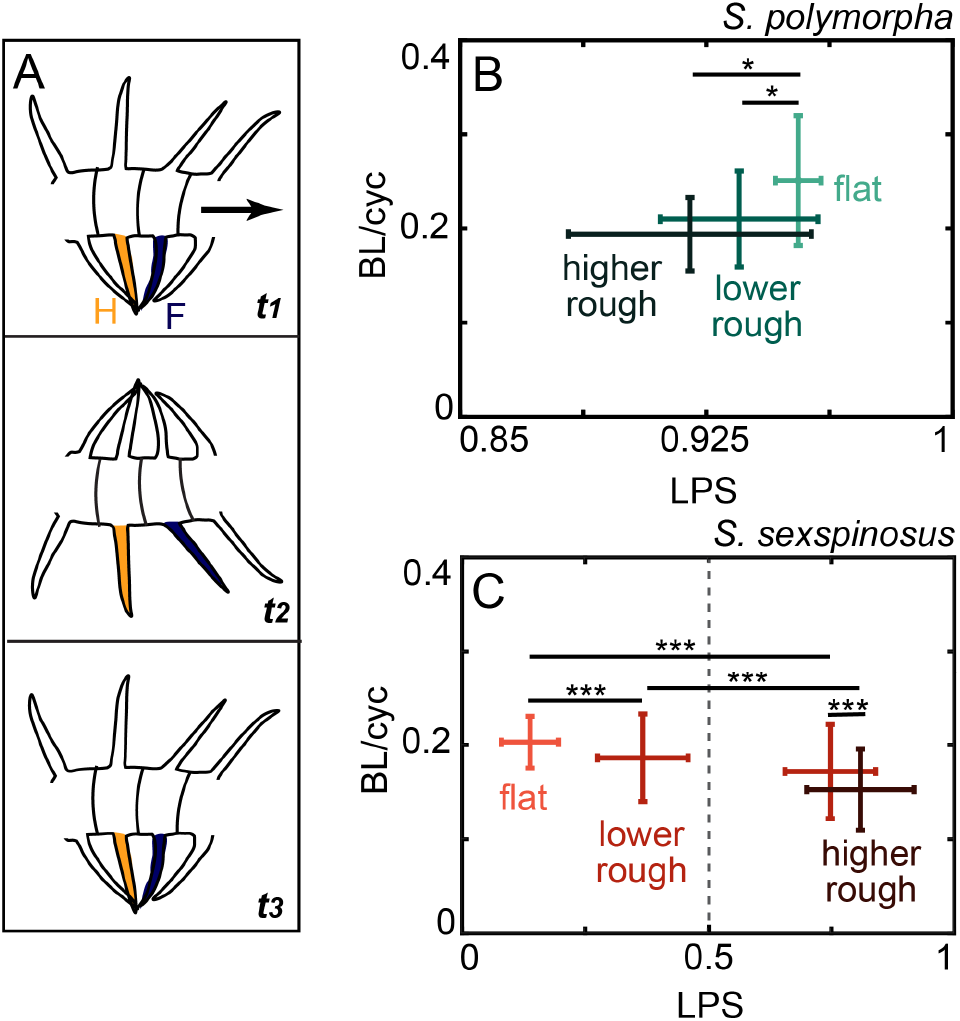
Centipede performance as a function of leg phase shift. (A) Diagram of centipede segment, moving over time. Arrow shows direction of motion (left to right). Blue limb denotes forelimb, *F*, and yellow limb denotes adjacent hindlimb, *H*. Leg phase shift (LPS) corresponds to the fraction of the time a hindlimb moves in the same direction as the adjacent forelimb. (B) Displacement per gait cycle of *S. polymorpha* as a function of LPS on each terrain. Light green, medium green, and dark green, correspond to flat, lower rough, and higher rough terrain, respectively. (C) Displacement per gait cycle of *S. sexspinosus* as a function of LPS on each terrain. Dashed lines corresponds to LPS = 0.5. LPS < 0.5 corresponds to direct limb-stepping patterns. LPS > 0.5 corresponds to retrograde limb-stepping pattern. Light orange, medium red, and dark red, correspond to flat, lower rough, and higher rough terrain, respectively. Differences were significant at p≤0.05, p≤0.001 for one and three asterisks, respectively. For *S. polymorpha*, LPS differences were significant at p≤0.05 between flat and higher rough terrain. For *S. sexspinosus* LPS differences were significant at p≤0.05 between flat and lower rough terrain (LPS<0.5), and at p≤0.001 between flat and lower rough terrain (LPS>0.5), between flat and lower rough terrain (LPS>0.5), between flat and higher rough terrain, between lower rough terrain with distinct LPS, and between lower (LPS<0.5) and higher rough terrain.

We calculated the speed of both centipedes species for all terrains and quantified the limb-stepping behavior by calculating the LPS. *S. polymorpha* achieved a speed of 0.19±0.04 body lengths per gait cycle (BL/cyc) and LPS of 0.92 ±0.04 on the flat terrain (Figure 5B). With increasing terrain complexity, there was a decrease in both speed and LPS. *S. polymorpha* achieved speeds of 0.21±0.05 and 0.25±0.07 BL/cyc and a LPS of 0.93±0.02 and 0.95±0.01, for the lower and higher rough terrain, respectively. Previous studies have characterized the relationship between speed and body undulation; when traveling at high speeds, centipedes that use a retrograde limb-stepping pattern display an increase in body undulation, specifically an increase in the maximum body wave amplitude (15). Conversely, when these centipedes travel at low speeds, there is a significant decrease in the maximum body wave amplitude making body undulation negligible (15). Thus, lack of body undulation observed in *S. polymorpha* on the rough terrains (Figure 3B(ii),B(iii)) may correspond to a decrease the speed. We note that although on flat terrain the centipedes displayed some body undulation, centipedes were not stimulated such that maximum speed was elicited on any of the terrains. In other words, we allowed centipedes to move at their preferred traveling speed.

*S. sexspinosus* achieved a speed of 0.20±0.03 BL/cyc, with a LPS of 0.14±0.06 on the flat terrain (Figure 5C), consistent with observations of the use of a direct limb-stepping pattern (15). On the lower rough terrain, trials were categorized by LPS. *S. sexspinosus* had wide distribution of LPS throughout each trial; interestingly, there are two clusters of LPS (one in the direct regime, the other in the retrograde regime) in the spectrum. When the centipede used only direct waves, it achieved a speed of 0.19±0.05 and used a LPS of 0.37±0.09. In contrast, the animal achieved a speed of 0.17± 0.05 and a LPS of 0.75±0.10 if the centipede used a retrograde wave. Unlike with a retrograde limb-stepping wave, previous studies have found that centipedes that use a direct wave do not use body undulation independent of speed. In the case of *S. sexspinosus*, when it used a retrograde wave it did not exhibit body undulation. While a retrograde limb-stepping pattern is related to body undulation, it remains unknown why the centipedes did not exhibit body undulation. Lack of body undulation can be due to: 1) the inability of the centipede to generate and propagate traveling waves of body curvature, or 2) higher speeds not being elicited. On higher rough terrain, *S. sexspinosus* used only retrograde limb-stepping waves; the centipede achieved a speed of 0.15±0.04 and a LPS of 0.81±0.11. A retrograde limb-stepping pattern facilitates “follow the leader” between limbs; when a single limb is placed on the ground, the rest of the limbs follow. Thus, we posit the centipede modulated the LPS to reduce the uncertainty of limb-substrate placement.

Phase over time for each leg (*φ*_*i*_) was calculated to find the retraction (*T*_*ret*_) and stride (*T*_*stride*_) period (Figure 6A-B). *T*_*ret*_ and *T*_*stride*_ is the time associated with backward movement of a limb during stance and the duration of a gait cycle, respectively (see Materials and Methods). These were used to calculate duty factor (*DF*), stride frequency (*ω*_*stride*_), step length (*L*_*step*_), and stride length (*L*_*stride*_) for both *S. polymorpha* and *S. sexspinosus* across all terrains (Figure 6C-F).

**Fig. 6.**
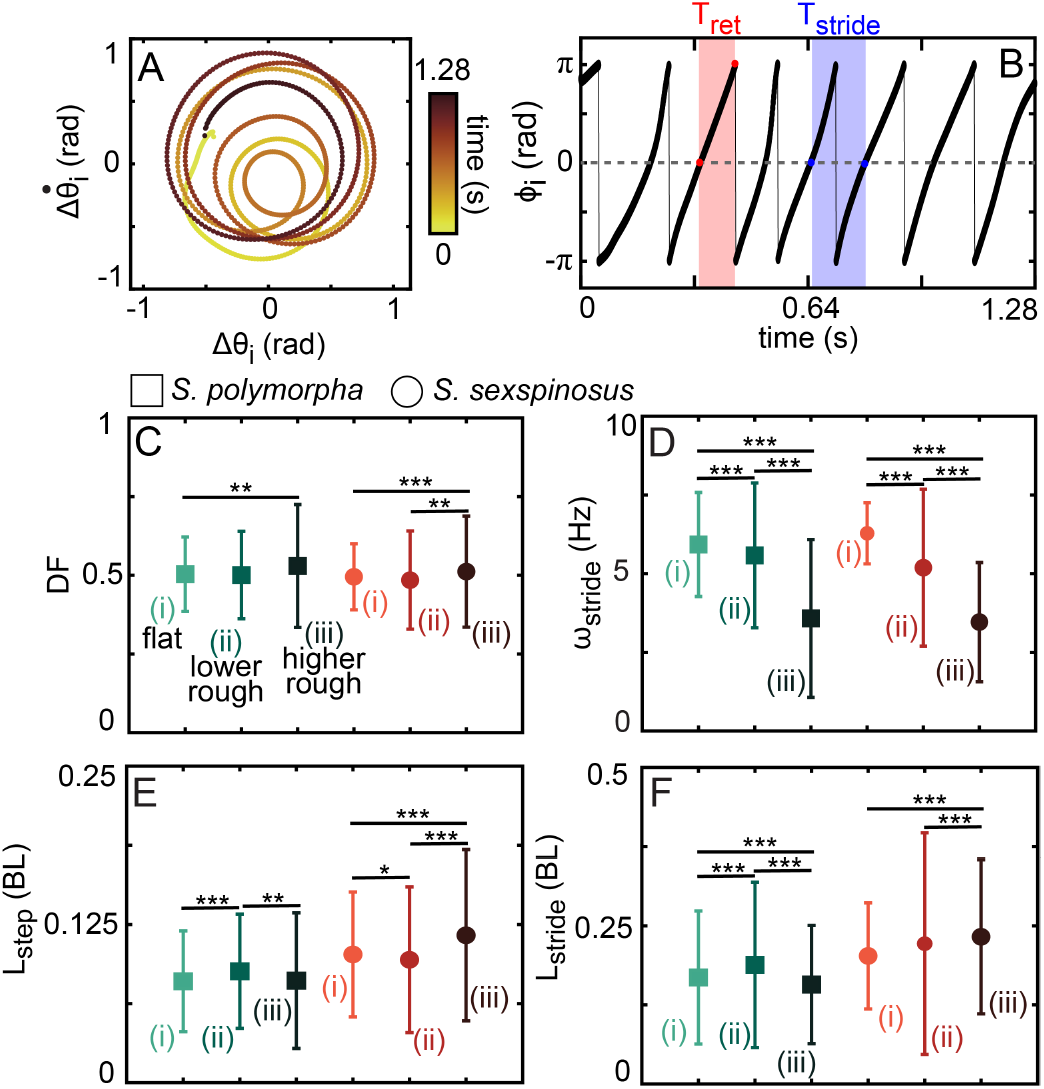
Centipede leg parameters. (A) Phase portrait for the angles (Δ*θ*_*i*_) of a single leg of *S. polymorpha* on flat terrain, colored by time, obtained from Hilbert transform. (B) Phase, *φ*_*i*_, of a single leg of *S. polymorpha* on flat terrain over time. Red band highlights a single retraction period, *T*_*ret*_. Blue band highlights a single stride period, *T*_*stride*_. (C) Duty factor, (D) stride frequency (*ω*_*stride*_), (E) step length (*L*_*step*_), and (F) stride length (*L*_*stride*_) for *S. polymorpha* and *S. sexspinosus* on all terrains. Green squares and orange circles correspond to *S. polymorpha* and *S. sexspinosus*, respectively. Numbering corresponds to (i) flat, (ii) lower rough, and (iii) higher rough terrain. Differences were significant at p≤0.05, p≤0.01, p≤0.001 for one, two and three asterisks, respectively.

Independent of terrain, both centipedes species achieved comparable *DF* (Figure 6C). *S. polymorpha* used a *DF* of 0.50± 0.12, 0.50±0.14,and 0.53±0.19, for flat, lower rough, and higher rough terrain respectively. *S. sexspinosus* used a *DF* of 0.50±0.11, 0.48± 0.16,and 0.51±0.18, for flat, lower rough, and higher rough terrain respectively. This suggests small modulations to the timing between the swing and the stance are sufficient to navigate these environments, potentially due to redundancy. In contrast, we observe a decrease in the stride frequency (*ω*_*stride*_) with increasing terrain complexity (Figure 6D). However, there was comparable *ω*_*stride*_ in each terrain between centipede species. *S. polymorpha* used a *ω*_*stride*_ of 6±1.7, 5.6±2.3, and 3.6±2.5 Hz, for flat, lower rough, and higher rough terrain respectively. *S. sexspinosus* used a *ω*_*stride*_ of 6.3± 1, 5.2±2.50,and 3.5±1.9 Hz, for flat, lower rough, and higher rough terrain respectively.

*L*_*step*_ and *L*_*stride*_ were leg parameters that could be directly impacted due to limb-substrate collisions. In other words, we expected the complexity of the rough terrains would cause the centipedes to modulate *L*_*step*_ or *L*_*stride*_. Surprisingly, *L*_*step*_ was not greatly affected independent of the terrain complexity (Figure 6E). *S. polymorpha* used a *L*_*step*_ of 0.08±0.04, 0.09±0.05, and 0.08±0.05 BL, for flat, lower rough, and higher rough terrain respectively. *S. sexspinosus* used a *L*_*step*_ of 0.10±0.05, 0.10±0.06,and 0.12±0.07 BL, for flat, lower rough, and higher rough terrain respectively. In the case of *S. sexspinosus*, the *L*_*step*_ remained relatively constant although these animals changed the direction of the limb-stepping pattern. Similarly, *L*_*stride*_ remained relatively constant independent of the terrain complexity for both centipede species (Figure 6F). *S. polymorpha* used a *L*_*stride*_ of 0.17±0.11, 19±0.13, and 0.16±0.09 BL, for flat, lower rough, and higher rough terrain respectively. *S. sexspinosus* used a *L*_*stride*_ of 0.20±0.08, 0.22±0.18,and 0.23±0.12 BL, for flat, lower rough, and higher rough terrain respectively. Although *S. sexspinosus* exhibits changes in the gait *L*_*step*_, and *L*_*stride*_ were averaged for all trials. Averages included when the animals were using both direct and retrograde limb-stepping patterns. Therefore, some of the variance could be attributed to intervals in which these centipedes switched from direct to retrograde.

### Passive limb mechanics

For both centipede species, we observed limb-substrate collisions along the horizontal plane due to the height disparities between adjacent blocks. Instead of jamming (i.e., limbs getting stuck/caught) into a block, the centipede’s limbs passively bent in the direction the force from the block was applied on the limbs (towards the body, opposite to the direction of motion) (SI Movie 2). We posit the inherent flexibility of the centipede’s limbs facilitated this passive limb behavior, which we have termed “passive gliding”. In previous studies (27), we found similar dynamics (i.e., passive mechanical compliance) improved locomotive performance on complex terrain of a centipede robot model without changes in the control. Thus, we hypothesize passive gliding allows centipede to negotiate limb-substrate collisions without actively modulating the limb-stepping pattern.

We identified each instance (when one or more limbs were bent due to a block at any point in time) of passive gliding for every trial of both centipede species. Figure 7A,C, show examples of passive gliding on the lower rough terrain for *S. polymorpha* and *S. sexspinosus*, respectively. We observed *S. sexspinosus* displayed greater instances (27 instances) of passive gliding than *S. polymorpha* (19 instances). However, the number of occurrences is path dependent; different paths led to different number of occurrences of passive gliding per individual (Figure 7B,D). A single *S. sexspinosus* did not exhibit any passive gliding, while all *S. polymorpha* exhibited passive gliding. Figure 7E,G, show examples of passive gliding on the higher rough terrain for both centipede species. *S. polymorpha* and *S. sexspinosus* displayed 53 and 60 instances of passive gliding, respectively. The number of limb-substrate collisions increased on the higher rough terrain due to the increase of complexity and height disparities between adjacent blocks. Unlike in the lower rough terrain, where few limbs collided with the block, a greater number of limbs would exhibit passive gliding on the higher rough terrain (Figure 7A,C). In addition, in some trials passive gliding occurred on both sides of the body. Figure 7C shows and example in which limbs and both side of the body engage in passive gliding simultaneously.

**Fig. 7.**
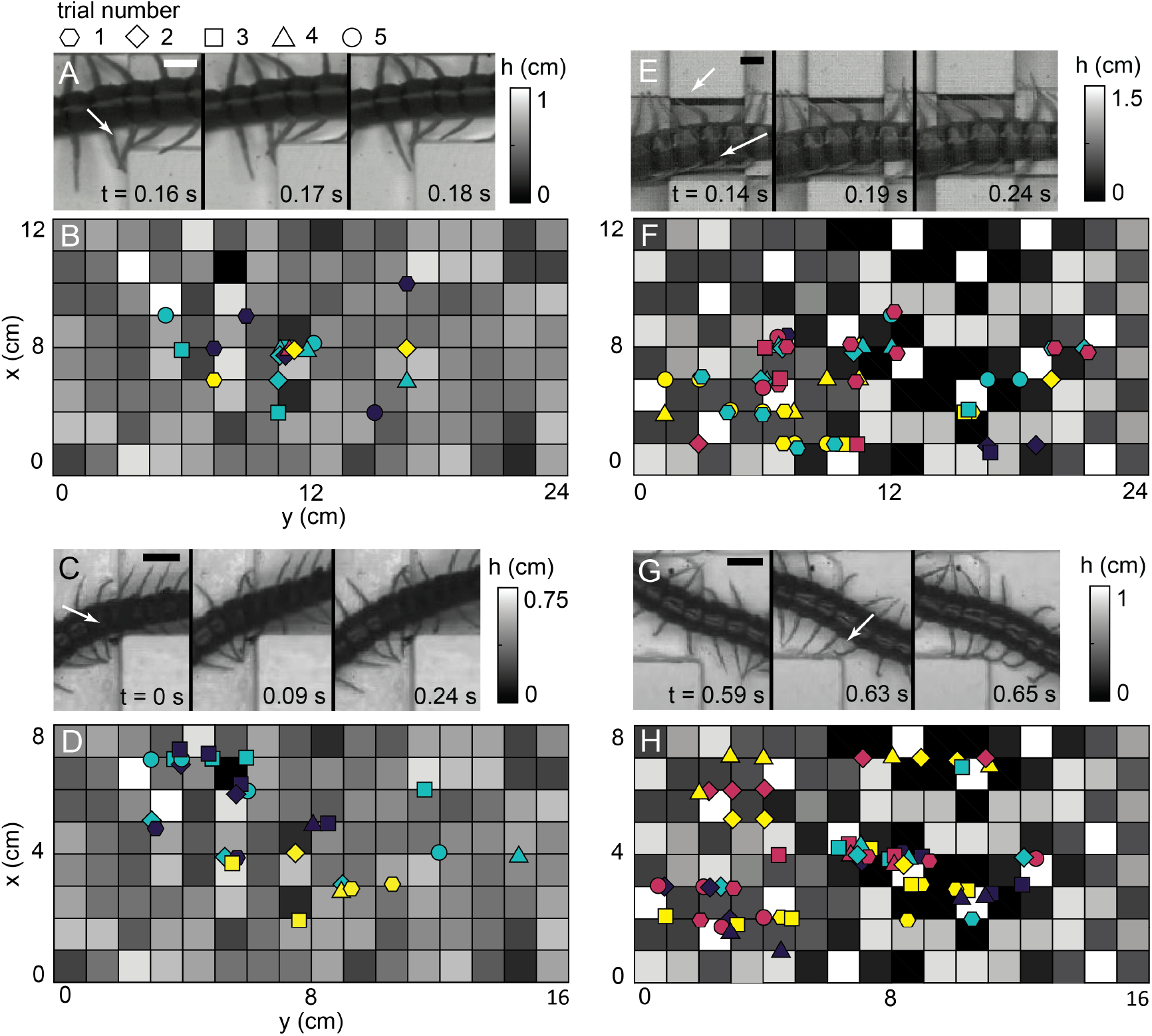
Passive limb gliding during obstacle interference. (A) Snapshots of passive limb gliding during limb-substrate collisions and (B) locations where horizontal limb-substrate collisions occurred for *S. polymorpha* on the lower rough terrain. Upon collision between a limb and a block, animals continued across the terrain without modulating their stepping patterns. Arrow highlights the limb-substrate interaction. (C) Snapshots of passive limb gliding during limb-substrate collisions and (D) locations where horizontal limb-substrate collisions occurred for *S. sexspinosus* on the lower rough terrain. (E) Snapshots of passive limb gliding during limb-substrate collisions and (F) locations where horizontal limb-substrate collisions occurred for *S. polymorpha* on the higher rough terrain. (G) Snapshots of passive limb gliding during limb-substrate collisions and (H) locations where horizontal limb-substrate collisions occurred for *S. sexspinosus* on the higher rough terrain. Distinct shapes and colors correspond to individual trials and animals, respectively. All scalebars correspond to 0.75 cm.

We calculated the probability density function (PDF) of body and limb angles for all terrains. Figure 8A(i)-(ii) show the PDF of body angles for *S. polymorpha* and *S. sexspinosus*, respectively. On flat terrain, PDF are centered at 0 for both centipede species. With increasing terrain complexity, the tails of the PDFs increase, corresponding to larger bends on the body. Neither of the centipede species exhibited body undulation on the rough terrains. However, because of the complexity of the terrains, there was a higher likelihood that centipedes would not travel in a straight path. Thus, bends on the body reflected in the distributions are related to movements of the centipedes when moving from one row of blocks (down or up the page) to another. PDF for legs on the left and right side of the body are shown in Figure 8B(i)-C(ii) for *S. polymorpha* and *S. sexspinosus*, respectively. With increasing terrain complexity, we observe an increase in the shift of the peak of the distributions. Due to the path dependence and passive gliding observed in the centipedes, PDFs obtained for the left and right side of the body are distinct. Thus, shifts from the peak are more prominent for the legs on the right side of the body, corresponding to limb-substrate collisions on the right side of the body.

**Fig. 8.**
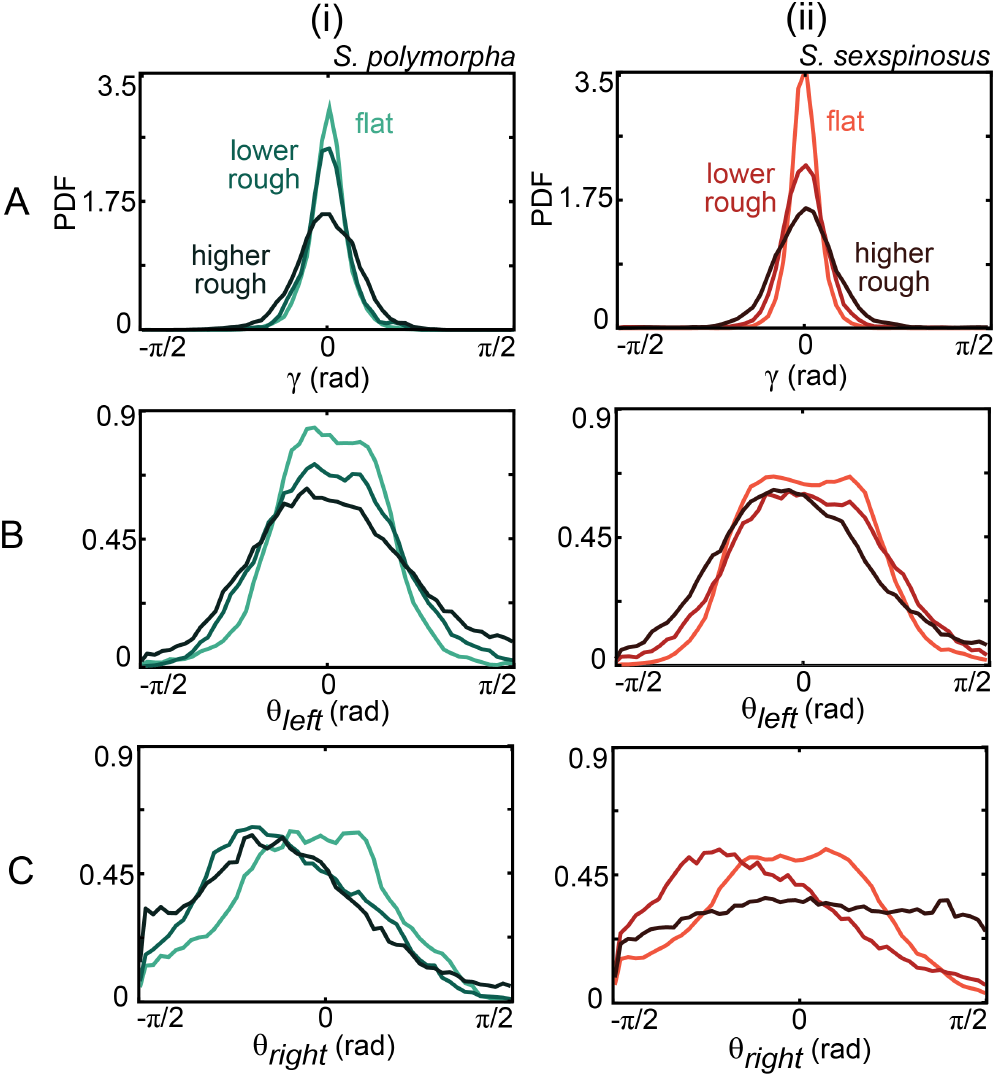
Probability of passive limb gliding across terrains. Probability density functions (PDF) of the (A) body, (B) left limbs, and (C) right limbs, for both (i) *S. polymorpha* and (ii) *S. sexspinosus*. For *S. polymorpha*, light green, medium green, and dark green, correspond to flat, lower rough, and higher rough terrain, respectively. For *S. sexspinosus*, light orange, medium red, and dark red, correspond to flat, lower rough, and higher rough terrain, respectively.

## DISCUSSION

Experiments revealed that the centipedes’ *DF, L*_*step*_, *L*_*stride*_ is not greatly affected, even in highly rugged terrains, potentially due to the redundancy in these animals (35). For example, redundancy in the limbs may increase robustness and facilitate terrain traversal, even when a limb is compromised (e.g., damaged or lost) (30). We observed a decrease in *ω*_*stride*_ with increasing terrain complexity. However, further investigation is necessary to understand the biomechanical advantage that these animals obtained with lower *ω*_*stride*_.

We found that *S. polymorpha* and *S. sexspinosus* leverage passive mechanics to navigate rough terrains. When a limb collided with an obstacle, the limb-stepping patterns was minimally perturbed and obstacle negotiation was facilitated by passive limb flexion. Instead of precisely controlling every degree of freedom associated with many limbs, the animals leveraged the inherent flexibility of their limbs. Offloading the control into the mechanics is an effective strategy seen in other biological (13) and has been successfully implemented in synthetic (robots) (27; 36; 14; 28) locomotors. Moreover, the effect of passive mechanical elements for multi-legged systems has been previously studied on myriapod robophysical models (27; 28). Flexible passive components facilitated complex terrain traversal of the robot, whereas rigid components achieved limited performance (27).

We posit these strategies (passive mechanics and redundancy) are advantageous in these centipede’s natural environments. *S. polymorpha* is a desert dwelling centipede *S. sexspinosus* can be found in forests within leaf litter, detritus, and under rotting logs. Both of these centipedes must contend with heterogeneities and height disparities inherent of the many materials in their surroundings. Further, these animals compete for resources (e.g., food) and thus reducing the energy required to negotiate complex terrains might be desirable.

It is commonly accepted that centipedes in the order Scolopendromorpha, Geophilomorpha, and Craterostigmorpha use retrograde limb-stepping waves, while those in the order Lithobiomorpha and Scutigeromorpha use direct limb-stepping waves (as characterized by Manton (16)). In addition, it is commonly accepted that the direction in which the limb-stepping pattern is propagated is fixed (one-species-one wave hypothesis (21)). *S. sexspinosus* exhibited behaviors contrary to both of these. While *S. sexspinosus* is of the order Scolopendromorpha (*Cryptopidae* family), instead of retrograde it uses direct limb-stepping waves on flat solid surfaces. Moreover, when locomoting on rough terrain, *S. sexspinosus* exhibited changes in the LPS corresponding to a change of limb-stepping pattern direction (i.e., change in gait). This change in LPS is not unique to *S. sexspinosus* or centipedes of the order Scolopendramorpha; previous studies have found change in the limb-stepping pattern with changes in substrate (18; 20; 21; 37) in different centipede species. Therefore, further investigation of centipede locomotion is necessary to evaluate and advance understanding across order. When encountered with a terrain with height disparities, *S. sexpinosus*, modulated the limb-stepping pattern accordingly. We observed comparable performance independent of the direction that the limb-stepping pattern was propagated (on the lower rough terrain). However, why this is a desirable strategy for this centipede remains unknown. We posit this increases the probability of finding a secure foothold. By using a retrograde limb-stepping pattern, the centipede can place a limb on the terrain and posterior limbs follow. Thus, the uncertainty associated with a direct limb-stepping pattern is reduced by switching to a retrograde pattern. Further investigation is necessary to understand the biomechanical advantage in this transition. Moreover, it is important to note that while *S. sexpinosus* used direct limb-stepping patterns on flat surfaces, that is not the centipede’s natural environment. Therefore, it is important consider that the limb-stepping patterns these centipedes use in nature may be retrograde and not direct.

We note that the kinematic analysis of these experiments were constrained to two dimensions (along the long and short axis of the centipede). However, movement along the height of surface (out/into the page) could play an important role. In flat terrain, these centipedes exhibit lifting of the body during locomotion (SI Movie 3). In rough terrains, these animals may fall into cavities formed by the blocks where lifting of the body to continue terrain traversal is essential. In other instances, centipedes have segments of the body suspended in air while crossing large gaps (SI Movie 3). At those instances, it is possible the limb dynamics may change (i.e., from periodic leg movement to no movement when crossing gaps (1; 32)) depending on the local surroundings for each body segment. Further examination is required to understand contributions in 3 dimensions.

## CONCLUSION

We performed, to the best of our knowledge, the first experiments with myriapods on terrains with features modeling natural habitats and terradynamic complexity. We explored the effects of terrain complexity on body and leg dynamics on two centipede species, *S. polymorpha* and *S. sexspinosus*. Both of the centipedes species studied were from distinct environments. Yet, these animals leveraged their morphology and physiology to traverse complex terrain. We observed these animals used passive gliding of the limbs during limb-substrate interaction, minimizing gait perturbation and facilitating traversal. Further, on rough terrains, *S. sexspinosus* exhibited changes in the LPS corresponding to a change in the direction the limb aggregates were propagated (i.e., from direct to retrograde propagation).

In *S. sexspinosus*, active changes in the gait may reflect the plasticity of these animals in response to changes in the environment. This may be due to selective pressures related to the variability of the composition of this centipede’s environment. Comparable locomotive performance between centipede species and previous robotic studies (27) suggests that changes in gait do not lead to improved performance. Thus, further investigation is required to explore what drives changes of gait and how these changes are advantageous to the animal.

Future comparative work could extend to other centipedes species to study their locomotive strategies in rough terrains. This could offer insight to the environmental information these animals use to select a gait. In addition, 3 dimensional kinematic analysis may provide insight into the observed lifting of the body during locomotion (SI Movie 3) and the effects during rough terrain traversal. Moreover, future work could explore the locomotive performance as a function of the number of leg pairs on rough terrains with for not only live animals but also robophysical models, useful as scientific models and in tasks such as search and rescue.

## Acknowledgements

We thank Cari Kickerson, Carl Anthony, Daniel Foley and Christine Foley for capturing live animals. We thank Joseph R. Mendelson III for servings as point of contact and providing animals for experiments. We thank Alexandra Carruthers for initial prototyping of experiments. We thanks Yasemin Ozkan-Aydin for insightful comments and discussion.

## Contribution

KD, EE, BC, and DIG designed research and wrote the paper. EE recorded *S. polymorpha* and *S. sexspinosus* kinematics on all substrates and performed tracking; DS designed and manufactured rough terrains. KD analyzed biological data.

## Data availability

All data that support the findings of this study are included in the article and the supplementary material.

## Supplementary

Supplementary material available online.

